# Zero-echo time imaging achieves whole brain activity mapping without ventral signal loss in mice

**DOI:** 10.1101/2024.09.19.613868

**Authors:** Ayako Imamura, Rikita Araki, Yukari Takahashi, Koichi Miyatake, Fusao Kato, Sakiko Honjoh, Tomokazu Tsurugizawa

## Abstract

Functional MRI (fMRI) is an important tool for investigating functional networks. However, the widely used fMRI with T2*-weighted imaging in rodents has the problem of signal lack in the lateral ventral area of forebrain including the amygdala, which is essential for not only emotion but also noxious pain. Here, we scouted the zero-echo time (ZTE) sequence, which is robust to magnetic susceptibility and motion-derived artifacts, to image activation in the whole brain including the amygdala following the noxious stimulation to the hind paw. ZTE exhibited higher spatial and temporal signal-to-noise ratios than conventional fMRI sequences. Electrical sensory stimulation of the hind paw evoked ZTE signal increase in the primary somatosensory cortex. Formalin injection into the hind paw evoked early and latent change of ZTE signals throughout the whole brain including the subregions of amygdala.

Furthermore, resting-state fMRI using ZTE demonstrated the functional connectivity, including that of the amygdala. These results indicate the feasibility of ZTE for whole brain fMRI, including the amygdala and we first show acute and latent activity in different subnuclei of the amygdala complex after nociceptive stimulation.

**Significance statement:** Functional MRI (fMRI) is an important tool for investigating functional networks. However, the widely used fMRI in rodents has the problem of signal lack in the lateral ventral area of forebrain including the amygdala, which is essential for not only emotion but also noxious pain. Here, we used zero-echo time (ZTE) sequence, which was robust to susceptibility artifacts, for functional imaging in the whole brain including amygdala. We demonstrated the feasibility and advantages of using ZTE to investigate neuronal activity in mice. Furthermore, we first showed acute and latent activation in different subnuclei of the amygdala complex as well as other regions related to pain after nociceptive stimulation in mice.

## Introduction

Functional magnetic resonance imaging (fMRI) is used for noninvasive neuroimaging of the whole brain in rodents and humans. However, T2*-weighted image with gradient-echo echo-planar imaging (GE-EPI), a widely used sequence in mouse fMRI studies, causes the researchers to suffer from the susceptibility artifact (1). This susceptibility artifact induces image distortion and signal loss. In particular, the lack of signals in the amygdala and piriform cortex due to air in the ear makes it challenging to investigate neuronal activity in these regions in rodents (2). Limbic brain areas, including the amygdala, are essential for emotional responses such as fear, anxiety, and depression (3–5). The amygdala is also an important brain center for pain. Nevertheless, neuronal activity in the amygdala has not been fully investigated using fMRI because of this critical pitfall.

Zero-echo time (ZTE) MRI is a novel imaging technique that utilizes fast readouts, non-selective volume excitation, and 3D radial center-out k-space encoding. Thus, it can capture signals from short-T2 tissues. Furthermore, ZTE provides a robust image against susceptibility issues at the frontiers between tissues and air (6). Recently, multiband sweep Imaging with Fourier Transformation (MB-SWIFT) (7, 8) and ZTE (9) sequences have shown a remarkable reduction in sensitivity to magnetic susceptibility artifacts in the brain imaging of rats and humans compared to the GE-EPI sequence. Notably, a recent study indicated that ZTE has the potential to detect neuronal activity (10).Visual and whisker stimulation evokes ZTE signal changes in several brain regions, including the somatosensory cortex’s visual cortex and barrel field (10). However, no study has used ZTE to map the time course of amygdala activity.

Chronic pain is a major health problem worldwide, and the mechanisms underlying chronic pain have been investigated in model mice (11). A study in rodents reported that several brain regions contribute to acute and chronic pain, including the sensory cortices, thalamus, cingulate cortex, parabrachial nuclei (PB), amygdala, and hippocampus (12). The formalin test in mice is a valid and reliable nociception model sensitive to various classes of analgesic drugs (13). It has been reported that local injection of formalin induces biphasic nocifensive behavior within one hour (13) underlying the drastic time-dependent changes in neural activity in the brain occur, for example, the expression of the activity-dependent molecular marker c-Fos in the brain pain network, including the central amygdala, followed by plastic changes in synaptic transmission (14, 15). Because of the difficulty in imaging the amygdala using fMRI, we previously used manganese-enhanced MRI (MEMRI) to investigate the neuronal activation evoked by nociceptive pain caused by formalin injection (16). The merit of MEMRI is its robustness against susceptibility artifacts. MEMRI showed increased neuronal activity in several regions, including the insular cortex, nucleus accumbens, primary/secondary somatosensory cortex, and amygdala 6-12 hours after formalin injection. However, because MEMRI detects the accumulation of Mn^2+^ proportional to neuronal activity, it is challenging to assess the time-dependent changes in neuronal activation, from immediately after formalin injection through the latent the pain process using MEMRI.

Here, we hypothesized that ZTE would enable the investigation of neuronal activity in the amygdala. First, we assessed the feasibility of the ZTE sequence for fMRI in mice using electrical stimuli applied to the hind paw. We then investigated the time course of ZTE signal changes in the whole brain following nociceptive stimulation of the hind paw with formalin.

We also investigated the resting-state functional connectivity of the amygdala using ZTE.

## Results

### Comparison of image quality of EPI and ZTE

We compared the image quality of GE-EPI, spin-echo (SE)-EPI, ZTE at 1 °(ZTE-1), and ZTE at 5 °(ZTE-5) in the same mice. We compared SE-EPI because, in addition to GE-EPI, SE-EPI has also been used for rodent fMRI (17). Two flip angles were used for ZTE because the contrast depends on the flip angle (18). The significant issue with T2 and T2*-weighted EPI, compared to multi-slice Rapid Acquisition with Relaxation Enhancement (RARE) images, was not only the signal loss in the amygdala, piriform cortex, and entorhinal cortex near the ear canal due to air, but also the distortion of the brain’s shape (Fig. 1A-1C). In comparison, the ZTE images exhibited no signal loss in these regions and less distortion (Fig. 1D and 1E). The contrast of ZTE depends on the flip angle, and blood vessels, ventricles, and brain tissues can be distinguished with ZTE-5, indicating better contrast than ZTE-1. The multiple slices of GE-EPI (Fig. S1A) and ZTE-5(Fig. S1B), which cover the prefrontal cortex to the cerebellum, show that the image of GE-EPI is entirely distorted, with signal loss not only in the amygdala but also in the cerebellum and pons (Fig. S1A). In contrast, ZTE-5 did not show a lack of signal nor distortion in the whole brain (Fig. S1B). Additionally, eyes (Bregma +3.56) and ear cavity (Bregma −1.96 and −3.08) were clearly observed in ZTE-5 (white arrows in Fig. S1B). To evaluate the image quality of these sequences, we compared the spatial signal-to-noise ratio (SNR) of these sequences in the cortex. The SNR values of ZTE-1 and ZTE-5 were significantly higher than those of GE-EPI and SE-EPI (Fig. S2A). The temporal SNR (tSNR) was then calculated to assess the stability of baseline signal quality over time (19). The tSNR maps revealed that the tSNRs of ZTE-1 and ZTE-5 in the whole brain were much higher than those of the GE-EPI and SE-EPI (Fig. S2B). Further analysis of the tSNR within the sensorimotor cortex ROI showed that the tSNR of ZTE-1 and ZTE-5 were significantly higher than those of GE-EPI and SE-EPI (Fig. S2C). These results indicate the feasibility of ZTE sequence for the resting state fMRI.

**Figure 1.**
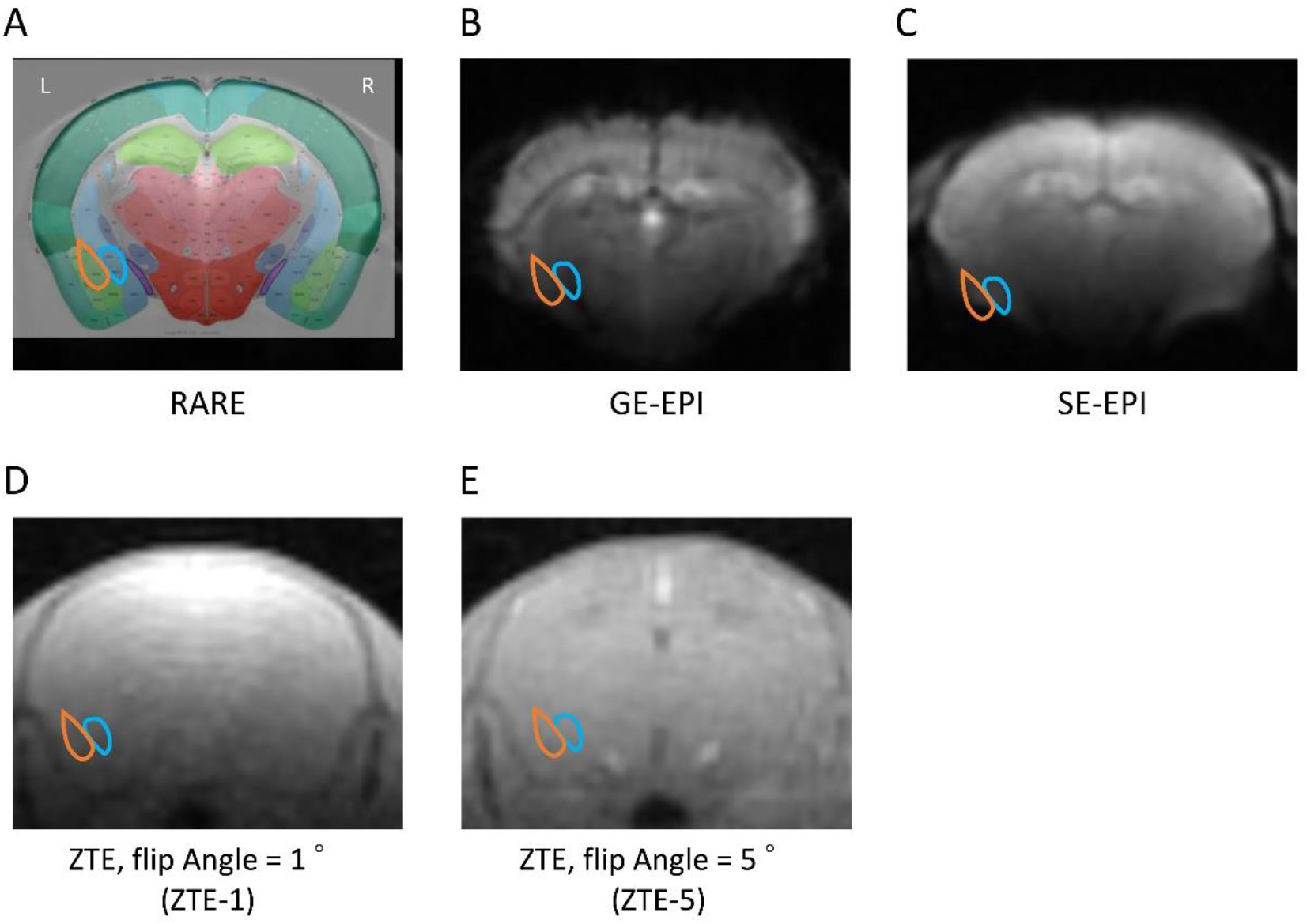
Images of EPI and ZTE. (A) Representative anatomical image by RARE. Orange and blue outlined nuclei indicate the LA/BLA and CeA, respectively, and the region is drawn with reference to the Allen Brain Atlas and Paxinos and Franklin’s The Mouse Brain in Stereotaxic Coordinates, with the locations of major structures landmarked (B-E). Representative images of GE-EPI (B), SE-EPI (C), ZTE, Flip Angle = 1 ° (ZTE-1) (D), and ZTE, Flip Angle = 5 ° (ZTE-5) (E).

### Signal changes by hind paw electrical stimuli

The response of ZTE signals to right hind paw electrical stimulation was compared with conventional fMRI (Fig. 2A). The time course of signal changes following hind paw electrical stimulation in the contralateral (left) primary somatosensory cortex was plotted. Hind paw stimulation induced positive signal changes in GE-EPI (Fig. 2B), ZTE-1 (Fig. 2C), and ZTE-5 (Fig. 2D). The percent changes in ZTE-1 were smaller than those in ZTE-5 and exhibited an earlier signal decrease after the end of stimulation compared to GE-EPI (Fig. 2E). The peak amplitude of ZTE-1 was smaller than that of ZTE-5 (1.21 ± 0.35 % for GE-EPI, 0.12 ± 0.04 % for ZTE-5, and 0.05 ± 0.02 % for ZTE-1), indicating that ZTE-1 has lower sensitivity to evoked neuronal activation than ZTE-5. The area under the curve (AUC) of the averaged time course showed that the AUC with ZTE-1 and ZTE-5 at Post2 (the period between 15 and 30 s after the end of stimulation) returned to baseline, while the AUC with GE-EPI at Post2 kept the increased signal to the baseline (pre) (Fig. 2F-2H). These results demonstrate the feasibility of using ZTE-5 and stimulation-induced fMRI. Based on these results, we decided to use ZTE-5 to image neuronal activation in the whole brain by intraplantar formalin injection.

**Figure 2.**
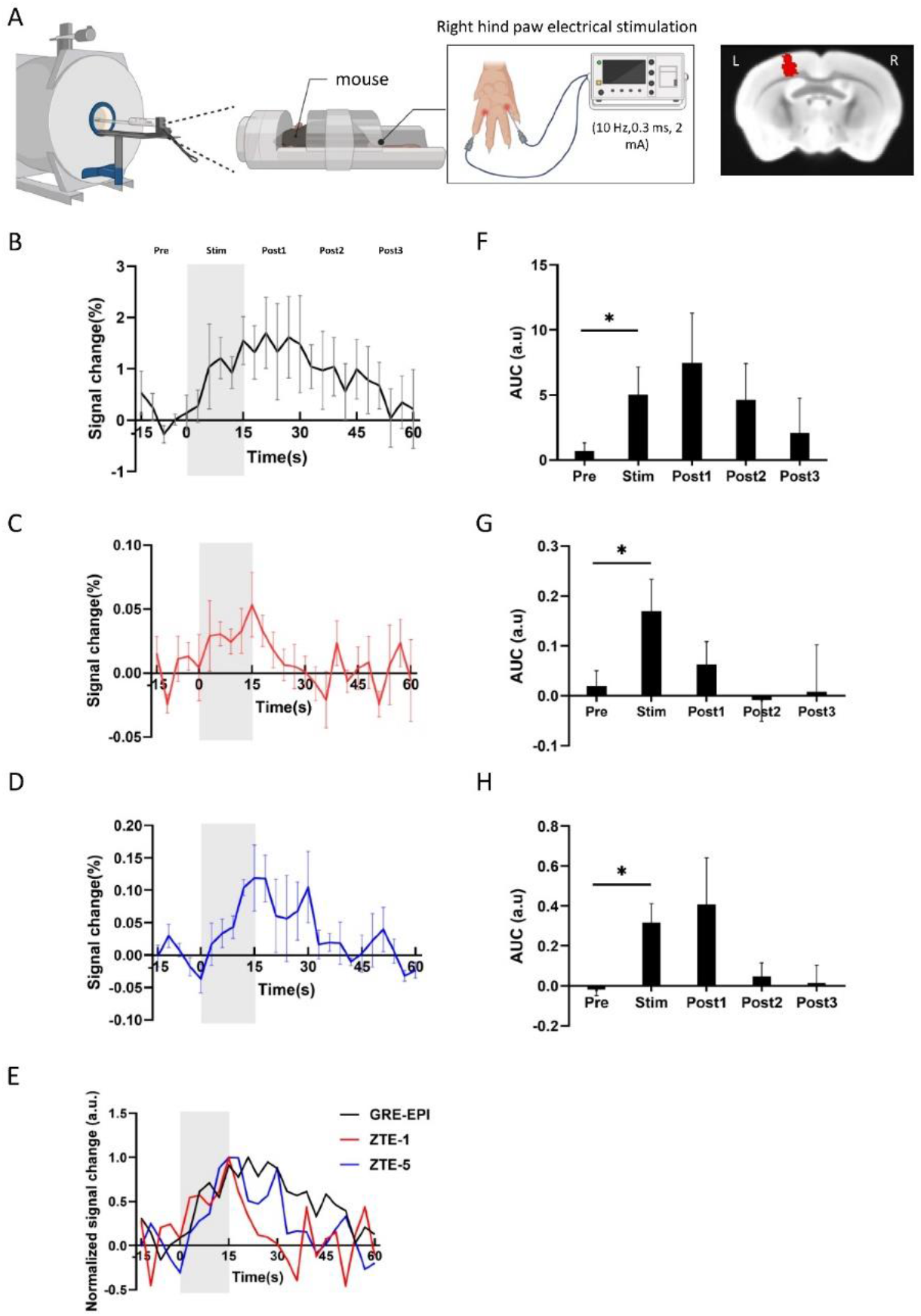
Time-course of primary somatosensory cortex evoked by hind paw electrical stimulation. (A) Experimental schematics (left) and the analyzed area of left primary somatosensory cortex (red on the right panel). (B-E) Averaged time-course of GE-EPI (B), ZTE-1 (C), and ZTE-5 (D) at the left primary cortex(S1HL). Grey zone indicates the right hind paw electrical stimulation. Data are expressed as mean ± S.E.M. (E) Normalized time-course of GE-EPI, ZTE-1, and ZTE-5 at the left primary sensory cortex of hindlimb. Grey zone indicates the right hind paw electrical stimulation. The peak of the averaged time course was normalized to 1. (F-H) Area under the curve (AUC) of averaged time-course of GE-EPI (F), ZTE-1 (G), and ZTE-5 (H). Post1, Post2, and Post3 represent the mean AUCs during the post-stimulation phases, specifically from 15 to 30 seconds, 30 to 45 seconds, and 45 to 60 seconds after the onset of electrical stimulation, respectively. Data are expressed as mean ± S.E.M. *p < 0.05 by paired t-test compared with Pre following one-way repeated measures ANOVA.

### Time-course of ZTE-5 signal changes in ROIs by intraplantar Formalin Injection

We visualized the time courses of ZTE-5 signal changes by acute formalin injection into the intraplantar surface of the right hind paw compared with saline in the whole brain, including the subregions of the amygdala (Fig. 3A and Table 1). Remarkably, biphasic signal changes were observed in several brain regions (Fig. 3B and 3C). The ZTE-5 signal increase was rapidly triggered by formalin injection (early surge). ZTE-5 signals in the contralateral (left) amygdala including the basolateral amygdala (BLA), central nucleus of the amygdala (CeA), and a part of bilateral basal ganglia circuit and ipsilateral (right) prefrontal cortex, increased following formalin injection. The signal increase in contralateral amygdala was prolonged until the end of scanning. The latent period of signal increase in the wide brain regions, including bilateral BLA, CeA, striatum, a part of hypothalamus and basal ganglia circuit, contralateral hippocampus, and contralateral motor cortex, occurred around 20 min (mid-term surge) after the injection (Fig. 3B and 3C). The signals in most regions of the bilateral thalamus, especially in the ipsilateral thalamus, decreased below the baseline following injection (Fig. 3B). We then investigated the correlation of the signal changes in typical brain regions related to pain (Fig. 3D). At 2.5 min after the injection, correlation of the ZTE-5 signal changes in several interhemispheric regions, such as ventromedial (VM) and mediodorsal (MD) thalamus, parabrachial nuclei (PB), BLA, and lateral amygdala (LA), was observed.

**Figure 3.**
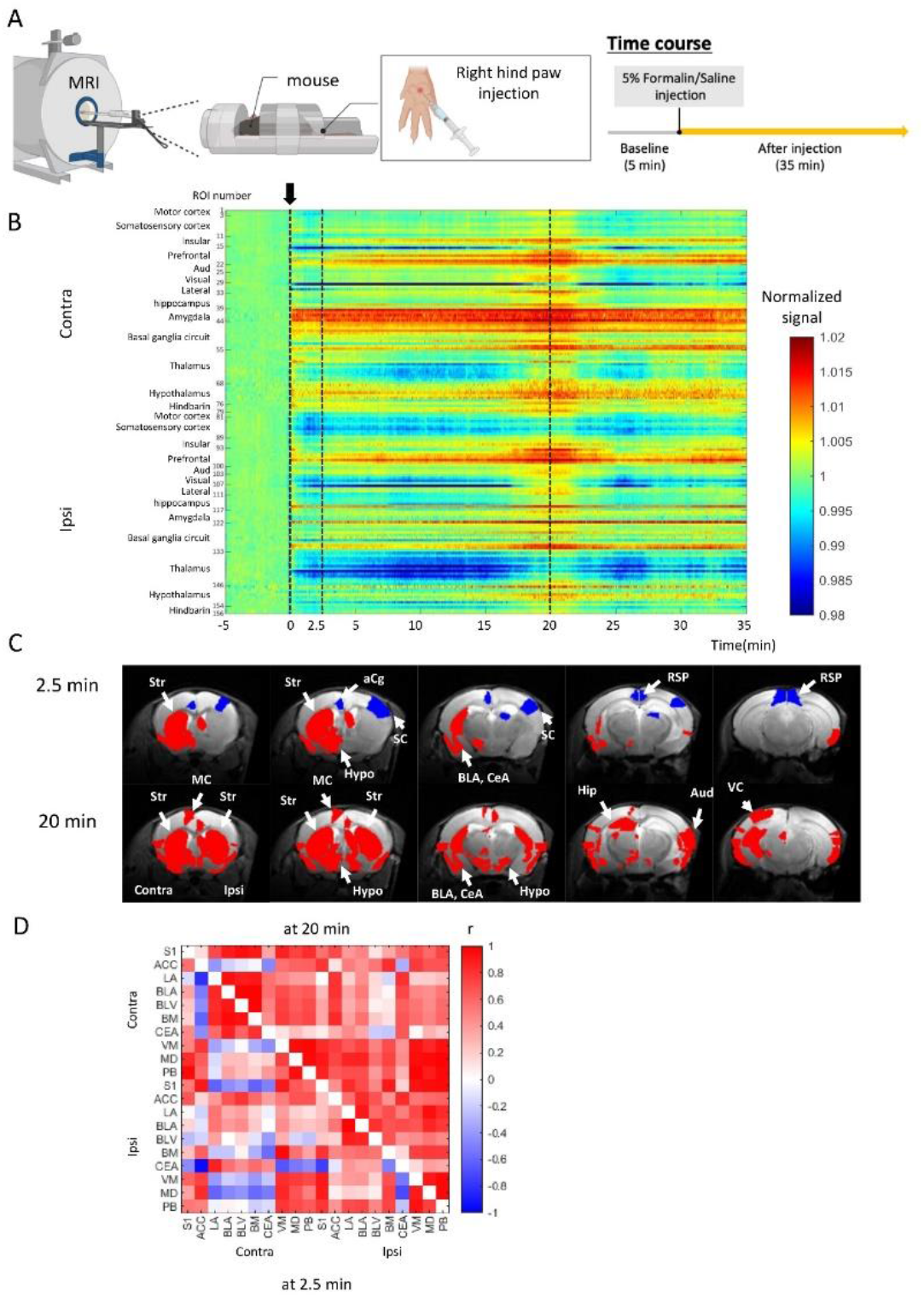
Time-course map of ZTE-5 signals in each ROI followed by the formalin injection. (A) Experimental schematics and formalin injection paradigms with ZTE-5. (B) Time-course map of averaged ZTE-5 signals in whole brain followed by the formalin injection. The color bar indicates the signal intensity normalized by the average of baseline intensity (-5 to 0 min) The horizontal axis indicates the time from the start of injection (*n* = 6). The left vertical axis indicates the brain regions of ipsi- and contra-lateral sides to the formalin injection. The numbers in vertical axis indicate the ROI numbers corresponding to Table 1. (C) Activation and deactivation maps. Red indicates signal increase (Normalized signal >1.005) and blue indicates signal decrease (Normalized signal < 0.99) compared to the baseline (-5 to 0 min). Aud, auditory cortex; BLA, basolateral amygdala; CeA, central amygdala; aCg, anterior cingulate cortex; Hip, hippocampus; Hypo, hypothalamus; MC, motor cortex; RSP, retrosplenial cortex; SC, somatosensory cortex; Str, striatum; VC, visual cortex. (D) Correlation matrices of normalized signal intensities between brain regions at 2.5 min (lower- left, *n* = 6) and at 20 min (upper-right, *n* = 6) after the formalin injection. Color bar represents the Pearson correlation coefficient between two brain regions. See Table 1 for the abbreviation.

**Table 1.**
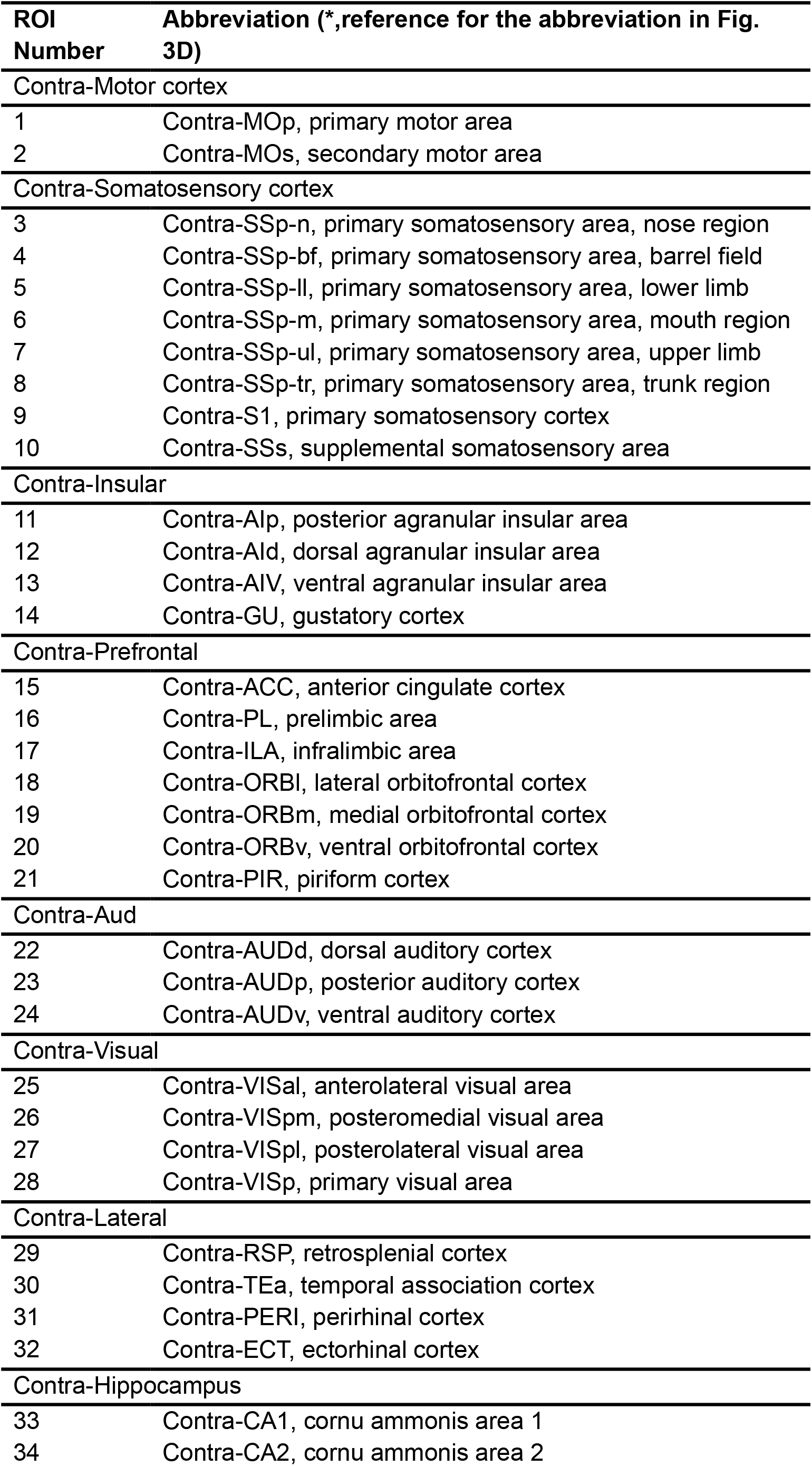
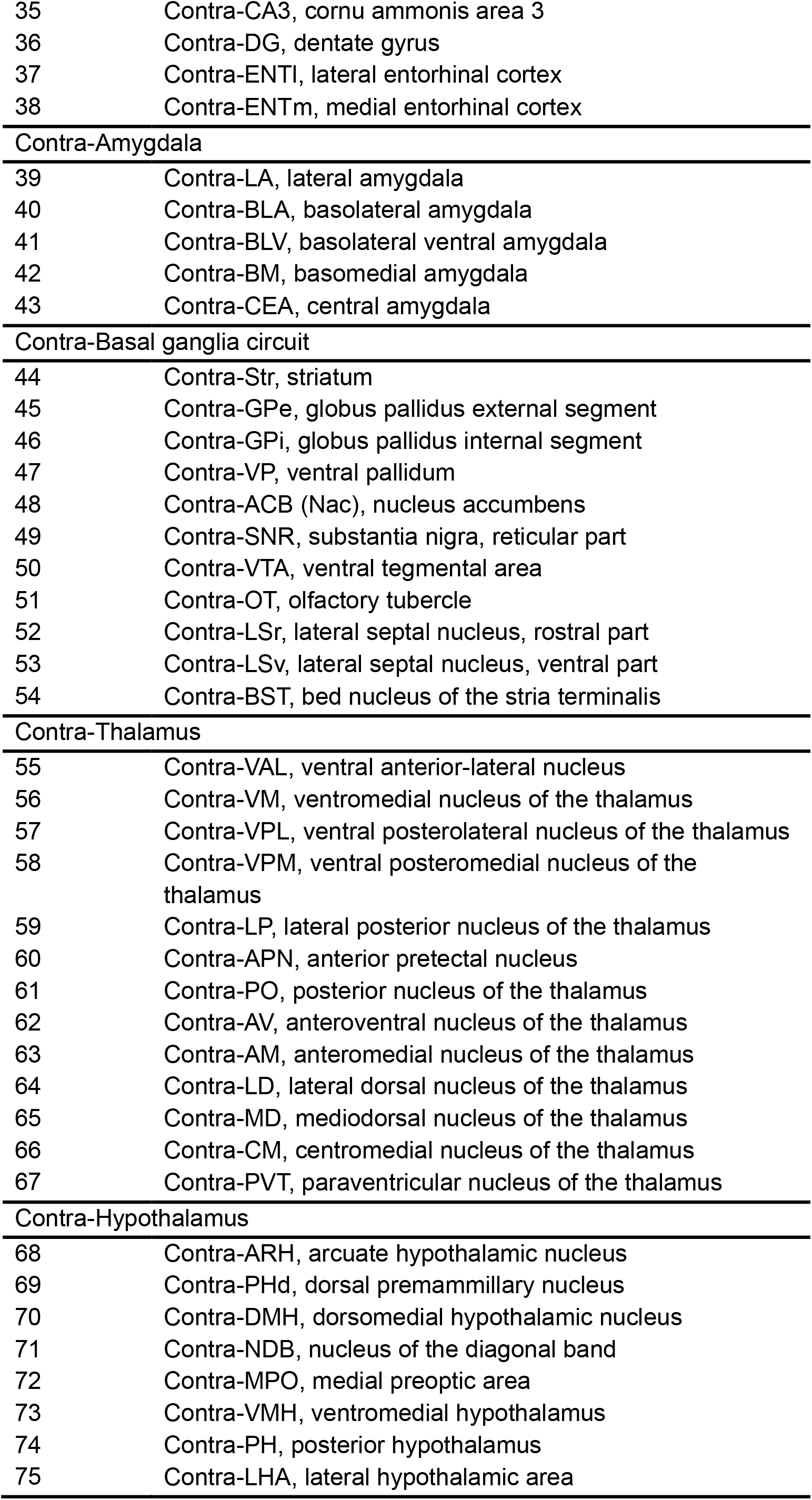
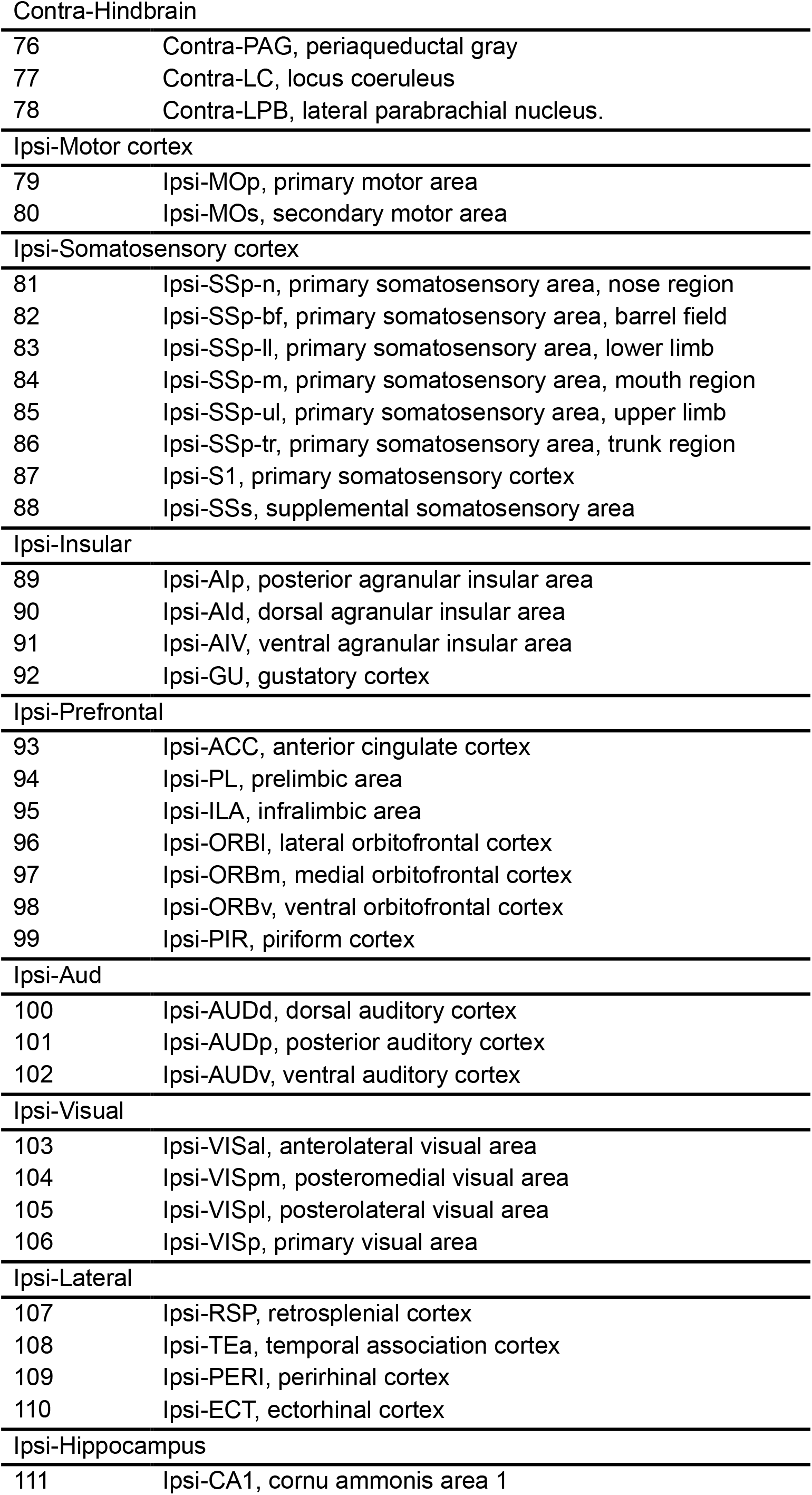
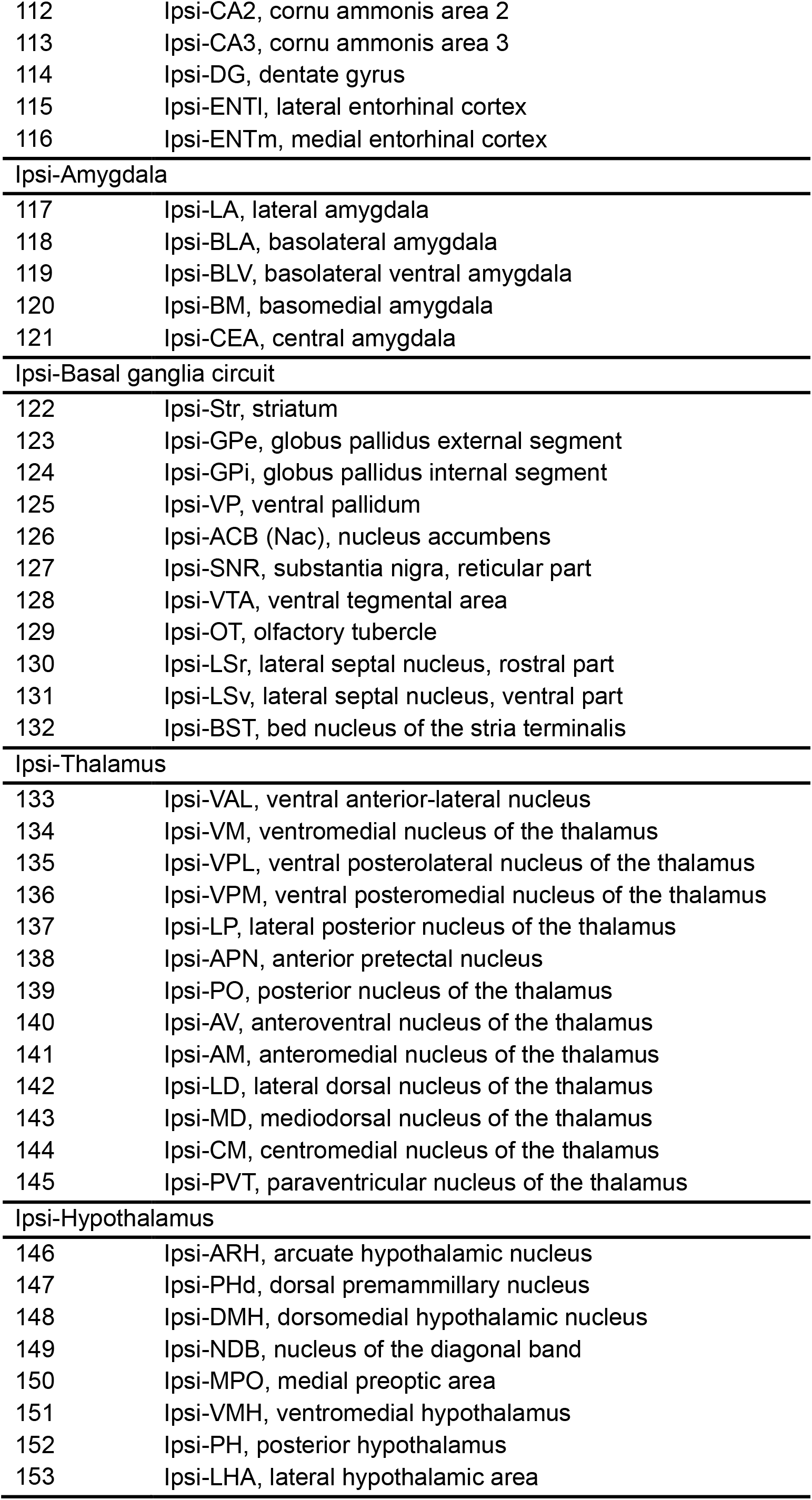
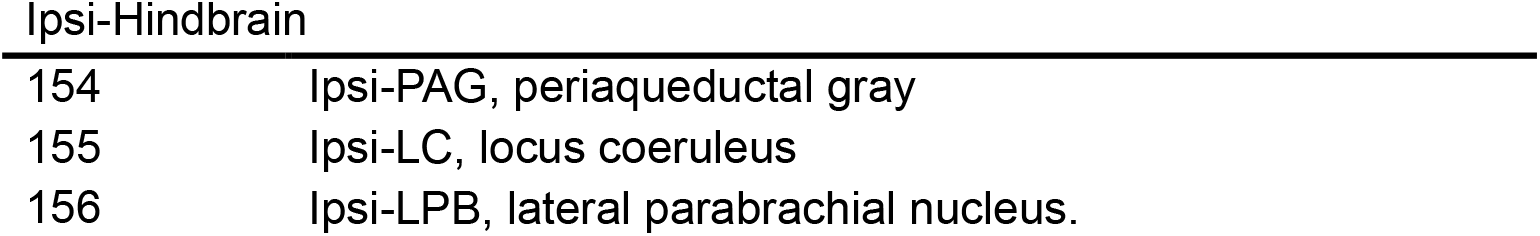
Regions-of-interest (ROIs) and abbreviations.

The negative correlation between contralateral (left) amygdala and ipsilateral (right) thalamus and S1 was also observed. In contrast, at 20 min after the injection, positive correlation was observed among wide regions except for the negative correlation between contralateral CeA and contralateral ACC (Fig. 3D). The time course of ZTE-5 signals in several brain regions related to pain is shown in Fig. 4. Significant signal increase was observed in the contralateral CeA (Contra-CeA), contralateral basolateral amygdala (Contra- BLA) and contralateral parabrachial nucleus (Contra-PB) (Fig. 4A, 4C, and 4E). Peak signal increases in Ipsi-CeA and Ipsi-BLA were observed during the latent surge (Fig. 4B and 4D). In contrast, no significant changes were observed in the ipsilateral PB (Ispi-PB), contralateral primary somatosensory cortex (Contra-S1) and bilateral primary visual cortex (Contra-V1 and Ipsi-V1) (Fig. 4F, 4G, 4I and 4J). The ZTE-5 signal in Ipsi-S1 was significantly decreased Immediately after injection only (Fig. 4H). A signal decrease except for 12-25 min was observed in the contralateral ACC (Contra-ACC) (Fig. 4K). Ipsilateral ACC (Ipsi-ACC) shows the transient signal increase after the injection and slight decrease around 10 min (Fig. 4L).

**Figure 4.**
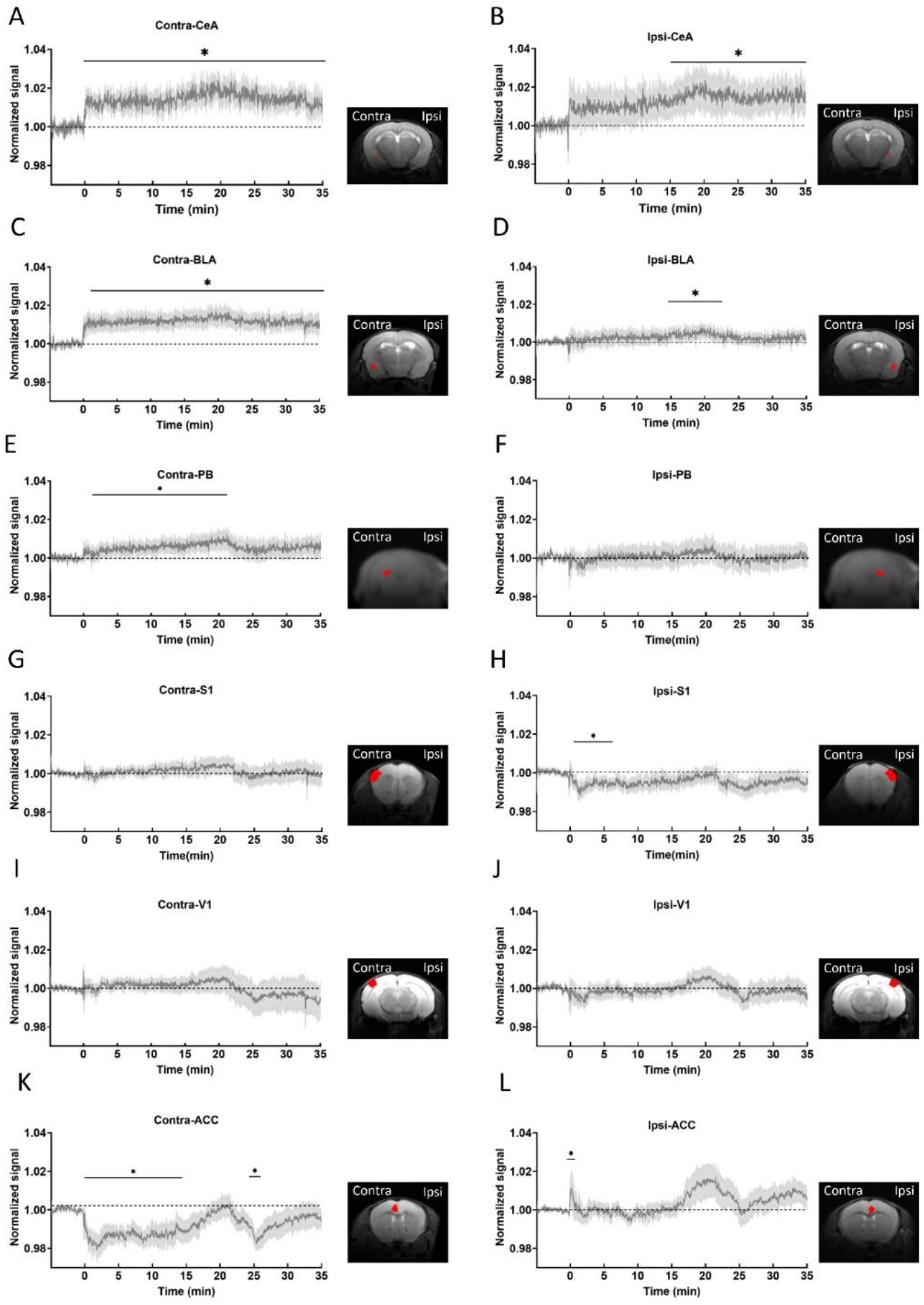
Time-course of ZTE-5 signals in the pain network regions following formalin injection. Averaged time-course of normalized signals in contra central nucleus of amygdala (Contra- CeA) (A), ipsi central nucleus of amygdala (Ipsi-CeA) (B), contra basolateral amygdala (Contra -BLA) (C), ipsi basolateral amygdala (Ipsi -BLA) (D), contra parabrachial nucleus (Contra -PB) (E), ipsi parabrachial nucleus (Ipsi -PB) (F), contra somatosensory cortex (Contra-S1) (G), ipsi somatosensory cortex (Ipsi-S1) (H), contra primary visual cortex (Contra -V1) (I), ipsi primary visual cortex (Ipsi -V1) (J), contra anterior cingulate cortex (Contra-ACC) (K), and ipsi anterior cingulate cortex (Ipsi-ACC) (L). Data are expressed as mean ± S.E.M. *p < 0.05 compared with averaged baseline following repeated measures ANOVA (n = 6).

Separated from the MRI experiment, we evaluated c-Fos expression, which is a marker of neuronal activation, in the bilateral CeA by formalin or saline injection (Fig. S3). Formalin injection induced the c-Fos expression bilaterally but c-Fos expression was not observed in the bilateral BLA. The saline injection did not induce c-Fos expression in these regions. This suggests that Fos is not equivalent to early neural activity.

### Functional connectivity by independent component analysis

It has been difficult to evaluate the functional connectivity of the whole brain including the subregions in the amygdala due to the difficulty of functional imaging in the amygdala (Fig. 1 and S1). Therefore, using ZTE, we examined resting state functional connectivity, including the amygdala. Functional connectivity was investigated in the resting state with GE-EPI, SE-EPI, ZTE-1, and ZTE-5 using independent component analysis (ICA). Regarding the functional connectivity including the amygdala, we found three components (C1 – C3) of unilateral functional connectivity, including left the amygdala, with ZTE-5 only (Fig. 5A).

**Figure 5.**
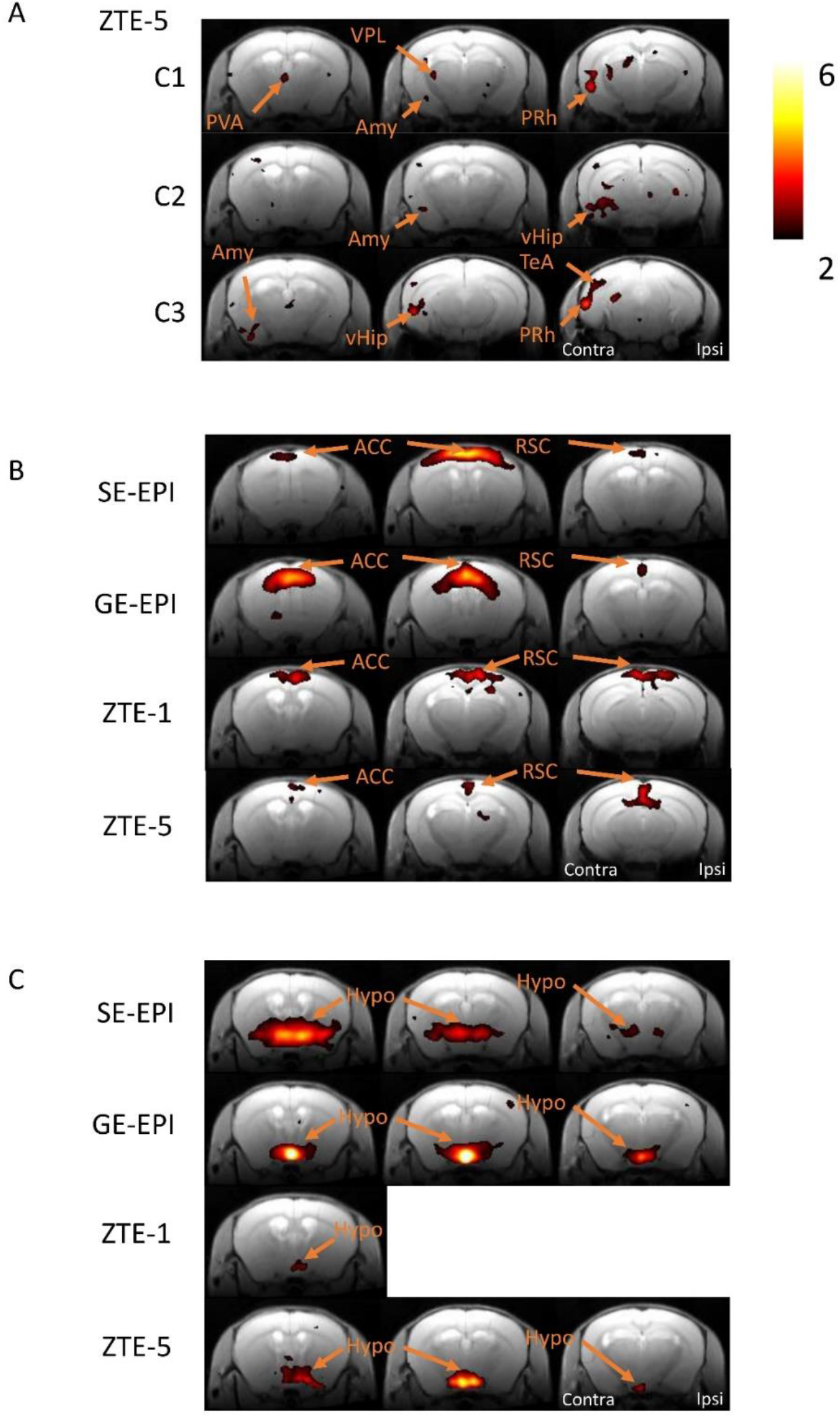
Functional connectivity. (A) Functional connectivity analysis of the whole brain, including the amygdala using the signals by ZTE-5 sequence and independent component analysis (ICA). Three components (C1, C2, C3) were extracted by independent component analysis with ZTE-5 only. (B) Default mode network with GE-EPI, SE-EPI, ZTE-1, and ZTE-5. (C) local network within hypothalamus with GE-EPI, SE-EPI, ZTE-1, and ZTE-5. Colar bar express the Z-values. ACC, anterior cingulate cortex; Amy, amygdala; Hypo, hypothalamus; PVA, paraventricular thalamus; PRh, Perirhinal cortex; RSC, retrosplenial cortex; vHIP, ventral hippocampus; VPL, ventrolateral thalamic nucleus.

These components are composed of left connectivity, including left primary visual area (PVA), left perirhinal cortex (PRh), ventral hippocampus (vHip), ventral posterolateral thalamic (VPL) and left temporal association cortex (TeA) (Fig. 5A). Importantly, typical functional connectivity, such as the default mode network (DMN), which is the anterior-posterior connectivity along the axis of the cingulate and retrosplenial cortices (Fig. 5B), and local connectivity, such as the hypothalamus, were observed in the GE-EPI, SE-EPI, ZTE-1, and ZTE-5 groups (Fig. 5C). These results indicate that ZTE-5 can detect functional connections like other sequences and can further extract functional connections including amygdala.

## Discussion

In the present study, we demonstrated the feasibility and advantages of using ZTE-5 to investigate neuronal activity in mice. Using this technique, we successfully demonstrated the neuronal activity in the amygdala. The critical issue with the commonly used fMRI sequence, that is, GE-EPI, in rodents is signal loss in the brain regions involving the amygdala and entorhinal cortex. However, fMRI of the amygdala is required to investigate its functional connectivity and neuronal activity because this region plays important roles in emotion, fear conditioning, and pain (20–22). ZTE-fMRI resolves this issue, and we demonstrated that the time course of whole-brain signals indicates biphasic activation in the formalin-induced inflammatory pain. This study demonstrated that ZTE-5 enables us to investigate the functional network, including the amygdala, in mice, similar to humans (23), and time-dependent dynamic brain activity immediately after the induction of inflammatory pain was unveiled. At 2.5 min, activities of many structures are oppositely affected (positively or negatively correlated depending on different areas), suggesting a localized increase in the ZTE signals, while those in were positively correlated in most of the regions at 20 min, suggesting that the surge in the brain activity at around 20 min after formalin injection occurred rather globally in the brain.

### The feasibility of ZTE-5 for fMRI

Previous studies have demonstrated the feasibility of using ZTE-5 for fMRI. Simultaneous electrophysiological recordings have shown that MB-SWIFT (24) detects neuronal activation in rats (8). In the present study, we compared tSNR, which affects false positives in fMRI (25), SNR, and hind paw electrical stimulation between two ZTE flip angles (1 ° and 5 °) and EPI images. The tSNR and SNR values of the ZTE-1 and ZTE-5 images were significantly higher than those of the GE-EPI and SE-EPI images (Fig. S1). Electrical stimulation of the hind paw revealed that ZTE-5 was more sensitive to neuronal activity. This is supported by a previous study in which the amplitude of the signal change with flip angles of 6° and 4° was higher than that with a flip angle of 2° (7). In this study, we used a maximum flip angle of 5° because of the power-output limitation of the RF amplifier. A higher flip angle should be assessed, depending on the conditions of the RF amplifier.

A previous study using group-level awake-state rat ICA results showed that MB-SWIFT fMRI could parcellate resting-state brain activity in awake rats (8). However, resting-state functional connectivity with ZTE in mice has not yet been conducted. Additionally, using the ZTE sequence, we showed functional connectivity involving the unilateral BLA, CeA, BMA, S1-bf, PRh, Ent, and TeA, in addition to the typical functional connectivity observed with EPI (Fig. 5). Anatomical projection exists between S1-bf and the BLA (26). Local circuit in the limbic system, including the BLA, CeA, ventral hippocampus and PRh have been studied using neuronal tracers (27). This indicates that ZTE-5 can detect the functional connectivity between the amygdala and other regions.

### Mechanism of ZTE-fMRI

We showed that the time course of the ZTE-fMRI signal change was different from that of GE-EPI during neuronal activation with hind paw electrical stimulation. What is the ZTE signal source that captures neuronal activity? Given that ZTE is not a T2* contrast, blood oxygenation level-dependent (BOLD) contrast (28) may not be a source of ZTE signal change. It is possible that blood inflow outside the excited volume still exists. A previous study applied a saturation pulse to MB-SWIFT, zero echo time and large bandwidth approach, to suppress signals from the inflow of blood during neuronal activation (7). The signal change was reduced by around 60% in the saturation band, but 40% of the signal change was still observed. This indicates that it does not rule out the possibility that other mechanisms may also play a role. A previous study has shown that astrocyte activation increases T1 relaxation (29). The mechanism of this phenomenon remains unknown; however, acute increases in T1 may reflect alterations in tissue microstructure, such as astrocyte activation and volume change. Astrocyte activation and volume changes are accompanied by neuronal activation to maintain extracellular substances, such as glutamate, potassium, and Ca^2+^ (30). Our previous studies have revealed that neuronal activation induces alterations in the extracellular space and restricts water diffusion, which is also influenced (25, 31). The earlier signal decrease after the end of somatosensory stimulation was similar to the earlier decrease in the diffusion-weighted MRI signals.

However, the onset time of ZTE was not different from that of GE-EPI. This indicated that the signal changes in diffusion-weighted fMRI and ZTE were derived from the same physical phenomenon. As the current study aimed to use ZTE for whole-brain mapping, including the subnuclei of the amygdala, the mechanism of ZTE, in addition to the inflow of blood, should be investigated in the future.

### Impact of amygdala imaging in neuroscience: Pain network

Pain is defined as “an unpleasant sensory and emotional experience associated with, or resembling that associated with, actual or potential tissue damage” (32). This definition implies that brain structures involved in any of these aspects of pain, such as nociception, sensation, and emotion, play essential roles in establishing the experience of pain. As multiple brain networks mediate the stimulation-pain link (33), revealing the dynamic temporal and spatial activation patterns in the brain network during the establishment of pain is crucial to understanding how acute nociception leads to more sustained suffering. This has been highly challenging with conventional functional brain imaging methods, mainly because of limited time resolution and limited capability of imaging the ventral structures, as discussed above. One such core brain structure linking nociception and pain is the amygdala. The amygdala, particularly the central subnucleus, receives direct nociceptive information via the parabrachial nucleus and sends information to widely distributed pain networks, such as the anterior cingulate cortex, insula, and prefrontal cortex, thereby playing a kernel role in the establishment of various aspects of pain (34–38).

The formalin-induced pain model is useful and widely used in preclinical studies of acute, inflammatory and latent sustained pain in rodents (15, 16, 39, 40). Subcutaneous injection of formalin triggers stereotypic behaviors consisting of early (phase 1) nocifensive behaviors starting immediately after injection, 2) a pause around 5-10 min and 3) late (phase 2) nocifensive behaviors starting around 15 min and lasting until ∼60 min post-formalin, each of which depends on distinct peripheral and spinal mechanism, such as TRP channel activation and central sensitization (39). An interesting observation in the present study was the ZTE-5 sequence-detected brain activities were composed of 1) early surge (2 to 15 min) accompanied by a decrease depending on the regions, 2) mid-term surge (around 20 min) accompanied by quiescence of the decrease, followed by moderate increase and decrease depending on regions. It would be of particular interest if these temporal changes in the ZTE- 5 signal reflect distinct activities of different brain regions receiving specific information from the nociceptive systems activated during each phase. If this is the case, such imaging with high spatiotemporal resolution would provide insights into how peripheral nociceptive events activate and suppress different brain networks involved in pain and suffering. To our surprise, the correlation between the two regions was also distinct at different time points (Fig. 3), indicating that the interrelationship between different pain-associated structures is not static but also shows dynamic transitions. Such minute-order analyses are difficult with the MEMRI approaches, which depend on sufficient intracellular accumulation of Mn2+ despite their advantage in capturing ventral structures near the inner ear (16). Altogether, these results underscore the potential of ZTE in neuroimaging studies involving the amygdala regions, where high temporal resolution is required to relate to behavioral changes.

## Materials and Methods

### Animals

Thirty-three C57BL/6J male mice (12-24 weeks) were allocated for the experiments: four for hind paw electrical stimulation, twelve for resting-state MRI, seven for formalin, and seven for saline injection, three for immunohistochemistry. Mice were maintained in a temperature-controlled (25 °C) environment on a 12h/12h light/dark cycle. All protocols were approved by the Institutional Animal Care and Use Committee of the Jikei Medical School of Medicine (Approval No. 2022-008) and the National Institute of Advanced Industrial Science and Technology (Approval No. 2024-0436). The animal experiment for immunostaining was approved by the Institutional Animal Care and Use Committee of the University of Tsukuba, and all procedures were conducted in accordance with the university’s Guidelines for Animal Experiments.

### Magnetic resonance imaging

All MRI experiments were conducted using a 9.4T scanner equipped with a cryoprobe (Bruker BioSpin, Ettlingen, Germany). The mice were first briefly anesthetized with 1.5 - 2 % isoflurane in the air for maintenance during head fixation. Isoflurane was decreased to 1.0 % to suppress the effect of the anesthetic on the BOLD signal when the respiratory rate was reduced to less than 100 breaths/min. Once the mouse head was fixed on the mouse bed, a bolus of medetomidine (0.1 mg/kg body weight, s.c.) was injected through the infusion cannula, which was inserted under the skin on the animals’ back. Then, it was infused continuously at 0.05 mg/kg body weight/h (s.c.) with 0.5%-1.0% isoflurane throughout the scanning. Body temperature was maintained by circulating hot water during the measurements. The MRI-compatible monitoring system (model 1025; SA Instruments, Stony Brook, NY, USA) measured the respiratory rate.

### MRI acquisition

The same parameters for MRI acquisition were used in all experiments. fMRI images with SE-EPI sequence were acquired with the following parameters: TR/TE = 3,000/20 ms, spatial resolution = 150 × 150 × 600 μm^3^ / voxel, 15 slices. fMRI images with GE-EPI sequence were acquired with the following parameters: TR/TE = 3,000/19 ms, spatial resolution = 150 × 150 × 500 μm^3^ / voxel, 15 slices. In ZTE acquisition, a 3D radial readout trajectory is used, and the free induction decay signal after excitation was recorded immediately. fMRI images with ZTE sequence were acquired with the following parameters: TR = 3,000 ms, flip angle = 1- or 5-degree, spatial resolution = 234 × 234 × 500 μm^3^ / voxel. Anatomical images were acquired for spatial correction using rapid acquisition with a relaxation enhancement (RARE) sequence following parameters: TR/effective TE = 2,500/33 ms, spatial resolution = 78 × 78 × 600 μm^3^ / voxel, RARE factor = 8, 4 averages.

### Resting state fMRI

Resting-state fMRI images were acquired from the same animal using GE-EPI, SE-EPI, ZTE-1, and ZTE-5. fMRI scanning was continued for 15 min in each sequence (300 volumes in total). The sequence of the MRI scans was randomly determined.

### Hind paw electrical stimulation

Two needle electrodes were inserted under the skin of the right hind paw, one between digits 1 and 2 and the other between digits 4 and 5. These electrodes were connected to an isolator (ISO-Flex, A.M.P.I., Israel) and a pulse stimulator (Master-8, A.M.P.I., Israel).

Rectangular pulses of 0.3 ms duration, 2 mA current, and 10 Hz were applied. The stimulation paradigm was a block design with six cycles of 15 s stimulus and 45 s post-stimulus periods.

### Intraplantar Formalin Injection

5% formalin solution (37% formaldehyde solution diluted in saline solution) or saline solution was acutely injected (100 μl) into the intraplanar surface of the right hind paw using a 24G catheter (Surflo I.V., Terumo, Japan) (41). Formalin or saline injection was injected 5 min after baseline imaging data collection, after which imaging data were acquired continuously for 35 min (total acquisition time was 40 min). Anatomical and functional MRI scans were acquired using the parameters above.

### fMRI analysis

#### Image preprocessing

The preprocessing of GE-EPI and SE-EPI, slice timing correction, spatial realignment, normalization, and smoothing were performed using SPM12. For ZTE, spatial realignment, normalization, and smoothing were performed using SPM12. For the resting-state fMRI analysis, further preprocessing was performed using the CONN toolbox (https://web.conn-toolbox.org/). The average signals in the cerebrospinal fluid, white matter, and head-motion parameters were used as nuisance regressors. Detrending and temporal filtering (0.008 – 0.08 Hz) were performed.

#### Signal-to-noise ratio (SNR) and temporal SNR (tSNR)

The SNR was calculated by dividing averaged signal intensity in ROI / standard deviation of the signal intensity in ROI of background.

The voxel-wise tSNR was calculated by dividing the temporal mean by the temporal standard deviation of the 300 volumes within each voxel.

#### Resting state fMRI analysis

Following the preprocessing of resting-state fMRI data, ICA was performed using the Group ICA of fMRI Toolbox (GIFT) software (https://trendscenter.org/software/gift/). First, all datasets underwent subject-specific principal component analysis (PCA), which estimated 150 components. All subjects’ reduced datasets were concatenated and underwent a PCA that estimated 30 components at the group level. Subsequently, a group-level spatial ICA was performed on the PCA output to identify 30 functional components.

#### Time-course of ZTE signal changes by formalin

Time-course analyses were conducted using a program written in MATLAB (MathWorks), as described previously (17, 25). The ROIs were registered using the Allen Brain Atlas (http://atlas.brain-map.org/). We then calculated the average signal values in the selected ROI using MATLAB. The percent changes in the ZTE-5 signals were calculated as follows:

Normalized signal intensity (a.u.) = (averaged signals within ROIs at each time point) / (averaged signals within ROIs during the basal period) The basal period was between 0 and 5 min before injection. The time-course signal changes to the baseline in formalin and saline injection groups were calculated by dividing the raw signals by averaged signal in the basal period at each time point. Because there was a signal drift during long time scanning, the signal change to the baseline after formalin injection at each time point was calculated as follows:

Normalized signal change to baseline in formalin group = (signal change in formalin group) / (averaged signal change across all subjects in saline group).

#### c-Fos immunohistochemistry

Separate from fMRI experiments, we examined the c-Fos expression to investigate the cellular level neuronal activation following formalin injection. Mice were anesthetized using the same method with fMRI and, after 90 min from the formalin or saline injection, transcardiac perfusion was performed. Mice were deeply anesthetized with 5% isoflurane in oxygen and intraperitoneally administered with Avertin. Then, mice were transcardially perfused with 0.1 M phosphate-buffered saline (PBS) containing heparin for 2 min, and then with ice-cold 4% paraformaldehyde (PFA) for 10 min. The brains were extracted and post-fixed overnight in PFA at 4°C, then transferred to 30% sucrose in PBS at 4°C for 2 days.

After incubation, the brains were embedded in Tissue-Tek O.C.T. Compound (Sakura) and stored at −80°C. Brains were sectioned with a cryostat (NX70, Thermo Scientific) in 50 µm thick sections. The sections were washed three times with PBS (pH 7.4), treated with 0.3% Triton X-100 in PBS for 5 min, and blocked in a solution containing 5% normal goat serum (NGS), 1% bovine serum albumin (BSA), and 0.3% Triton X-100 for 1 hour. The sections were then incubated overnight at 4 °C with rabbit anti-c-Fos antibody (1:2000, Abcam, ab190289) in the blocking solution. After washing three times with PBS, the sections were incubated with 5% NGS in PBS containing the secondary antibody (1:1000, Goat anti-Rabbit IgG (H+L) Highly Cross-Adsorbed, Alexa Fluor 488, Thermo Fisher, A11034) for 2 h at room temperature. The sections were then washed three times with PBS, mounted onto glass slides, and coverslipped with Mowiol (Sigma-Aldrich, St. Louis, MO). Brightfield microscopy Images were acquired using ZEISS Axio Zoom.V16.

## Statistics

Significant changes in the time course of MRI signals by formalin stimulation were evaluated using a paired t-test with averaged baseline (0-5 min before injection) following one-way repeated-measures ANOVA. The significance of AUC in electrical stimulation was assessed by paired-test with pre-period following one-way repeated-measures ANOVA. The significance of differences between more than three groups in SNR and tSNR was assessed using a t-test followed by Bonferroni correction.

## Acknowledgements

We appreciate Ms. Aki Omori for the support of this study.

---

**Figure S1.**
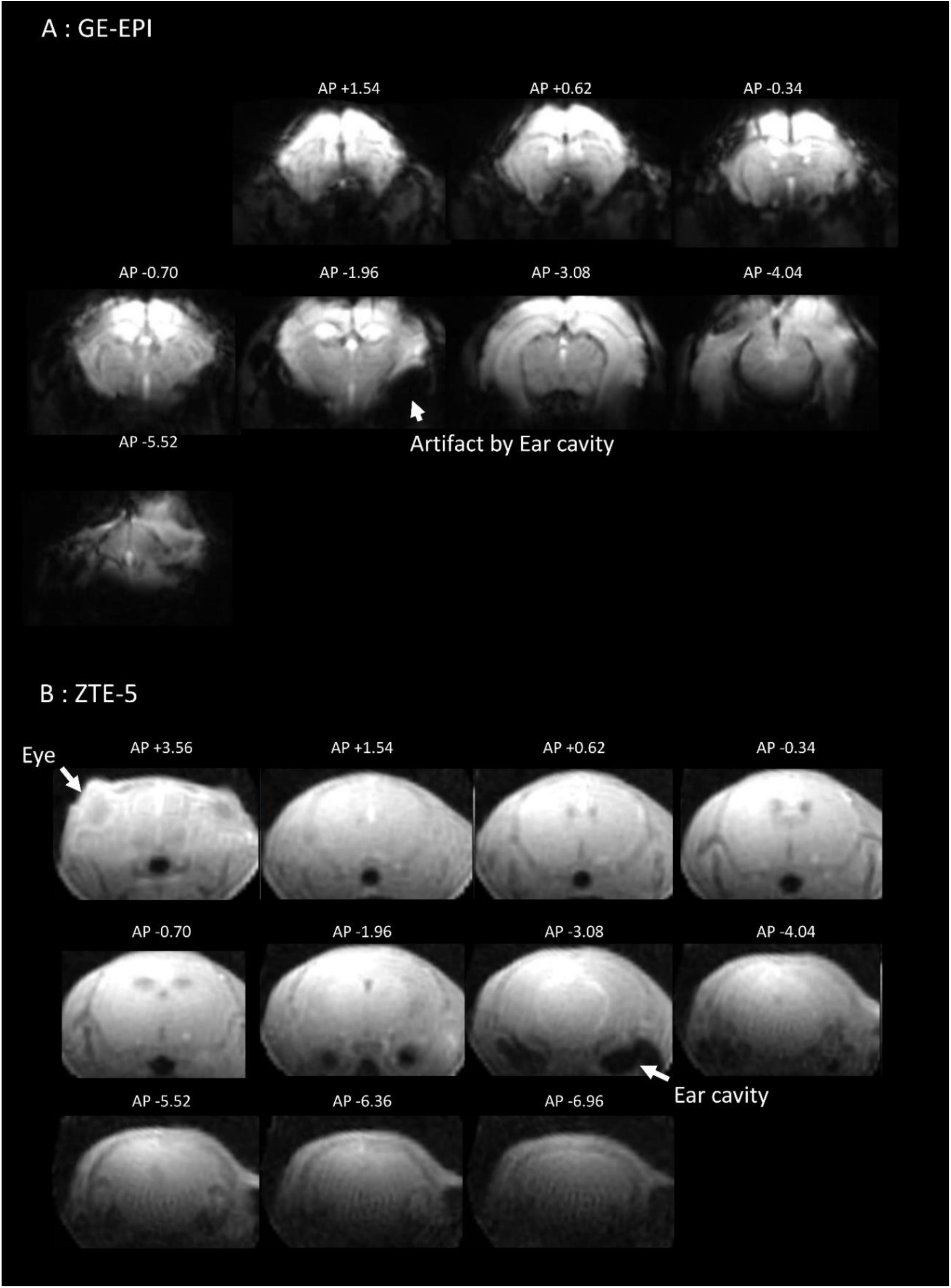
Representative images of GE-EPI and ZTE-5. Representative whole-brain images of GE-EPI (A) and ZTE-5 (B) from the same mouse.

**Figure S2.**
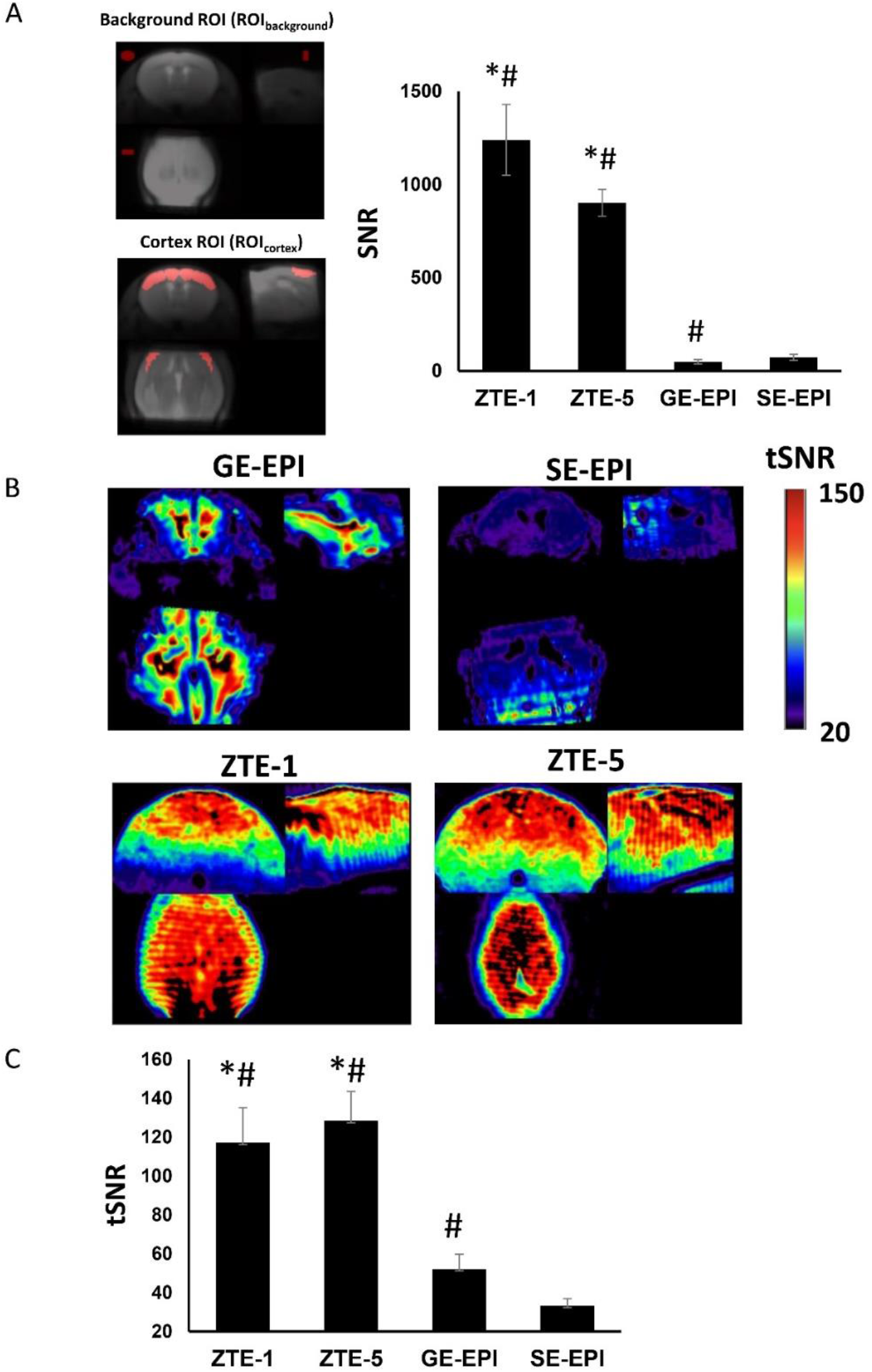
SNR and tSNR. (A) Left, each red area (top left: coronal, bottom left: horizontal, top right: sagittal) represents the background and cortex ROIs used for evaluating SNR in each imaging sequence. Right, SNR of the cortex ROI with ZTE-1, ZTE-5, GE-EPI, and SE-EPI. Data are expressed as mean ± SEM. *p<0.05, vs GE-EPI, ^#^p<0.05 vs SE-EPI by t-test with Bonferroni-correction (n=12). (B) Representative tSNR image of GE-EPI, SE-EPI, ZTE-1, and ZTE-5. Colar bar expresses the tSNR. (C) Averaged tSNR within sensorimotor cortex ROI. *p<0.05 vs GE- EPI, ^#^p<0.05 vs SE-EPI by t-test with Bonferroni-correction. Data are expressed as mean ± SEM.

**Figure S3.**
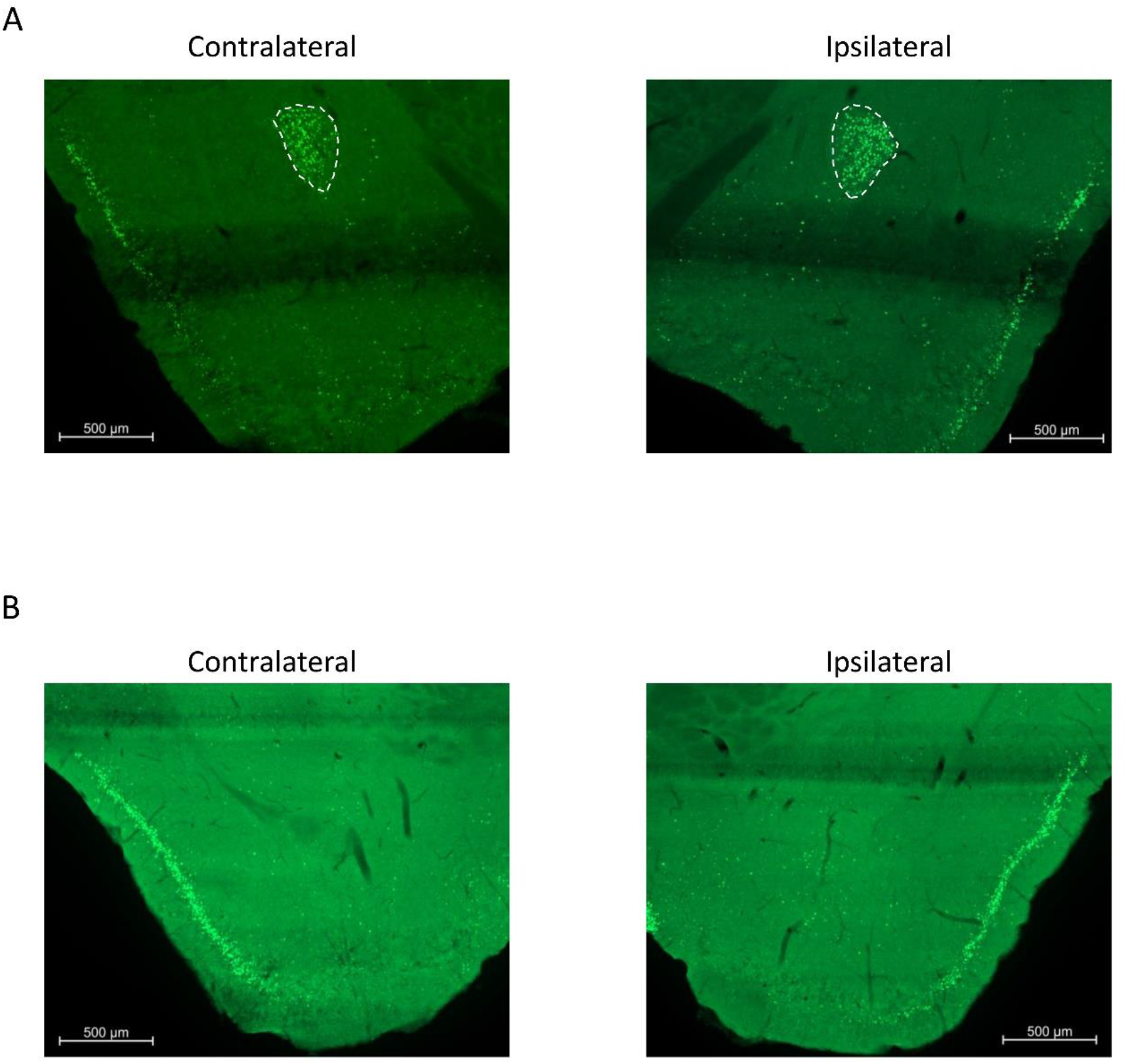
c-Fos expression following the formalin and saline injection into right hindpaw. Representative images of c-Fos protein expression in the bilateral amygdala at 90 min after formalin (A) /saline (B) injection into the right hind paw. The left and right panels are the images of the contralateral (left) and ipsilateral (right) sides of the formalin/saline injection. The areas outlined by dotted lines in the upper panels indicate the regions of strong c-Fos expression in the capsular and lateral parts of the central amygdala.

## References

1. M.-H. In et al., Correction of metal-induced susceptibility artifacts for functional MRI during deep brain stimulation. Neuroimage 158, 26–36 (2017).

2. R. Li et al., Restoring susceptibility induced MRI signal loss in rat brain at 9.4 T: A step towards whole brain functional connectivity imaging. PLoS One 10, e0119450 (2015).

3. K. J. Ressler, Amygdala activity, fear, and anxiety: modulation by stress. Biol. Psychiatry 67, 1117–1119 (2010).

4. V. Neugebauer, Amygdala pain mechanisms. Handb. Exp. Pharmacol. 227, 261–284 (2015).

5. J. Ferri et al., Blunted amygdala activity is associated with depression severity in treatment-resistant depression. Cogn. Affect. Behav. Neurosci. 17, 1221–1231 (2017).

6. C. Li, J. F. Magland, A. C. Seifert, F. W. Wehrli, Correction of excitation profile in Zero Echo Time (ZTE) imaging using quadratic phase-modulated RF pulse excitation and iterative reconstruction. IEEE Trans. Med. Imaging 33, 961–969 (2014).

7. L. J. Lehto et al., MB-SWIFT functional MRI during deep brain stimulation in rats. Neuroimage 159, 443–448 (2017).

8. J. Paasonen et al., Multi-band SWIFT enables quiet and artefact-free EEG-fMRI and awake fMRI studies in rat. Neuroimage 206, 116338 (2020).

9. E. Ljungberg et al., Silent zero TE MR neuroimaging: Current state-of-the-art and future directions. Prog. Nucl. Magn. Reson. Spectrosc. 123, 73–93 (2021).

10. P. Stenroos et al., How absence seizures impair sensory perception: Insights from awake fMRI and simulation studies in rats. eLife 10.7554/elife.90318 (2023).

11. C. X. Cui et al., Research progress on the mechanism of chronic neuropathic pain. IBRO Neurosci Rep 14, 80–85 (2023).

12. J. T. Da Silva, D. A. Seminowicz, Neuroimaging of pain in animal models: a review of recent literature. Pain Rep 4, e732 (2019).

13. S. Hunskaar, K. Hole, The formalin test in mice: dissociation between inflammatory and non-inflammatory pain. Pain 30, 103–114 (1987).

14. Y. Miyazawa, Y. Takahashi, A. M. Watabe, F. Kato, Predominant synaptic potentiation and activation in the right central amygdala are independent of bilateral parabrachial activation in the hemilateral trigeminal inflammatory pain model of rats. Mol Pain 14, 1744806918807102 (2018).

15. K. Shinohara et al., Essential role of endogenous calcitonin gene-related peptide in pain-associated plasticity in the central amygdala. Eur J Neurosci 46, 2149–2160 (2017).

16. D. Arimura et al., Primary Role of the Amygdala in Spontaneous Inflammatory Pain- Associated Activation of Pain Networks - A Chemogenetic Manganese-Enhanced MRI Approach. Front. Neural Circuits 13, 58 (2019).

17. T. Tsurugizawa, L. Ciobanu, D. Le Bihan, Water diffusion in brain cortex closely tracks underlying neuronal activity. Proc. Natl. Acad. Sci. U. S. A. 110, 11636–11641 (2013).

18. F. Wiesinger, M. L. Ho, Zero-TE MRI: principles and applications in the head and neck. Br J Radiol 95, 20220059 (2022).

19. K. Yoshida et al., Physiological effects of a habituation procedure for functional MRI in awake mice using a cryogenic radiofrequency probe. J Neurosci Methods 274, 38–48 (2016).

20. Y. Sun, H. Gooch, P. Sah, Fear conditioning and the basolateral amygdala. F1000Res. 9 (2020).

21. L. J. Becker et al., The basolateral amygdala-anterior cingulate pathway contributes to depression-like behaviors and comorbidity with chronic pain behaviors in male mice. Nat. Commun. 14, 2198 (2023).

22. C. Nicolas et al., Linking emotional valence and anxiety in a mouse insula-amygdala circuit. Nat. Commun. 14, 5073 (2023).

23. A. K. Roy et al., Functional connectivity of the human amygdala using resting state fMRI. Neuroimage 45, 614–626 (2009).

24. D. Idiyatullin, C. Corum, J.-Y. Park, M. Garwood, Fast and quiet MRI using a swept radiofrequency. J. Magn. Reson. 181, 342–349 (2006).

25. J. E. Perthen, M. Bydder, K. Restom, T. T. Liu, SNR and functional sensitivity of BOLD and perfusion-based fMRI using arterial spin labeling with spiral SENSE at 3 T. Magn. Reson. Imaging 26, 513–522 (2008).

26. I. M. Zakiewicz, J. G. Bjaalie, T. B. Leergaard, Brain-wide map of efferent projections from rat barrel cortex. Front. Neuroinform. 8, 5 (2014).

27. R. Kajiwara, I. Takashima, Y. Mimura, M. P. Witter, T. Iijima, Amygdala input promotes spread of excitatory neural activity from perirhinal cortex to the entorhinal- hippocampal circuit. J. Neurophysiol. 89, 2176–2184 (2003).

28. S. Ogawa, T. M. Lee, A. R. Kay, D. W. Tank, Brain magnetic resonance imaging with contrast dependent on blood oxygenation. Proc. Natl. Acad. Sci. U. S. A. 87, 9868–9872 (1990).

29. N. R. Sibson et al., Acute astrocyte activation in brain detected by MRI: new insights into T(1) hypointensity. J. Cereb. Blood Flow Metab. 28, 621–632 (2008).

30. M. Armbruster et al., Neuronal activity drives pathway-specific depolarization of peripheral astrocyte processes. Nat. Neurosci. 25, 607–616 (2022).

31. Y. Abe, K. Van Nguyen, T. Tsurugizawa, L. Ciobanu, D. Le Bihan, Modulation of water diffusion by activation-induced neural cell swelling in Aplysia Californica. Sci. Rep. 7, 6178 (2017).

32. S. N. Raja et al., The revised International Association for the Study of Pain definition of pain: concepts, challenges, and compromises. Pain 161, 1976–1982 (2020).

33. S. Geuter et al., Multiple Brain Networks Mediating Stimulus-Pain Relationships in Humans. Cereb Cortex 30, 4204–4219 (2020).

34. H. Bastuji, M. Frot, C. Perchet, M. Magnin, L. Garcia-Larrea, Pain networks from the inside: Spatiotemporal analysis of brain responses leading from nociception to conscious perception. Hum Brain Mapp 37, 4301–4315 (2016).

35. G. Corder et al., An amygdalar neural ensemble that encodes the unpleasantness of pain. Science 363, 276–281 (2019).

36. F. Kato, Y. K. Sugimura, Y. Takahashi, Pain-Associated Neural Plasticity in the Parabrachial to Central Amygdala Circuit : Pain Changes the Brain, and the Brain Changes the Pain. Adv Exp Med Biol 1099, 157–166 (2018).

37. T. Kiritoshi et al., Cells and circuits for amygdala neuroplasticity in the transition to chronic pain. Cell Rep 43, 114669 (2024).

38. E. Vachon-Presseau et al., The Emotional Brain as a Predictor and Amplifier of Chronic Pain. J Dent Res 95, 605–612 (2016).

39. C. R. McNamara et al., TRPA1 mediates formalin-induced pain. Proc Natl Acad Sci U S A 104, 13525–13530 (2007).

40. M. Miyazawa, A. R. Bogdan, K. Hashimoto, Y. Tsuji, Regulation of transferrin receptor-1 mRNA by the interplay between IRE-binding proteins and miR-7/miR-141 in the 3’-IRE stem-loops. RNA 24, 468–479 (2018).

41. X.-C. Fan et al., Hypersensitivity of Prelimbic Cortex Neurons Contributes to Aggravated Nociceptive Responses in Rats With Experience of Chronic Inflammatory Pain. Front. Mol. Neurosci. 11, 85 (2018).

